# The non-classic psychedelic muscimol suppresses inflammatory signaling and promotes neuroplasticity in schizophrenia-derived human cortical spheroids and astroglia

**DOI:** 10.64898/2026.04.08.717305

**Authors:** Ibrahim A. Akkouh, Jordi Requena Osete, Thor Ueland, Nils Eiel Steen, Ole A. Andreassen, Srdjan Djurovic, Attila Szabo

## Abstract

Schizophrenia (SCZ) is increasingly linked to neuroimmune dysregulation and impaired synaptic plasticity, yet the cellular mechanisms connecting inflammatory signaling to neural dysfunction remain poorly understood. Using human induced pluripotent stem cell (iPSC)-derived cortical spheroids (hCS) and astrocytes from patients with SCZ and matched controls, we investigated the effects of GABA_A_ receptor modulation on immune signaling and neuroplasticity. Inflammatory stimulation induced robust interferon-responsive transcriptional programs, prominently involving the antiviral effector MX1 and related interferon-stimulated genes. Computational deconvolution and cell type-specific analyses identified astrocytes as key mediators of these responses. Muscimol, a non-classic psychedelic and GABA_A_ receptor agonist, suppressed inflammatory gene expression, reduced secretion of proinflammatory cytokines, and attenuated interferon-associated signaling. In addition, muscimol induced neuroplasticity-associated transcriptional programs, including upregulation of *NTRK2* and *ELK1* in hCSs, and restored impaired glutamate uptake in iPSC-derived SCZ astrocytes. These effects were blocked by GABA_A_ receptor inhibition, confirming receptor-dependent mechanisms. Proteomic analyses of hCS cultures, and independent human dorsolateral prefrontal cortex datasets revealed baseline dysregulation of GABAergic and neurotrophin signaling in SCZ, supporting translational relevance. Together, these findings demonstrate that GABA_A_ receptor activation by muscimol suppresses inflammatory signaling while promoting neuroplasticity in hCSs, and identify astrocytes as central regulators of interferon-dependent neuroimmune dysfunction in SCZ. These results establish non-classic psychedelic compounds as potential modulators of neuroimmune-plasticity coupling and suggest that targeting astrocyte GABAergic signaling may represent a therapeutic strategy for restoring neural homeostasis in SCZ.

## INTRODUCTION

Schizophrenia (SCZ) is a severe neuropsychiatric disorder characterized by positive symptoms, such as hallucinations and delusions, as well as persistent cognitive deficits, negative symptoms and functional decline (1). Beyond its psychotic manifestations, SCZ is increasingly recognized as a disorder involving impaired neuroplasticity and aberrant brain maturation, contributing to long-term cognitive dysfunction and, in some patients, treatment resistance (2). Converging evidence from genetic, transcriptomic, and biomarker studies implicates immune dysregulation and low-grade neuroinflammation in the pathophysiology of SCZ, particularly in a biologically defined subgroup of patients exhibiting persistent symptoms despite antipsychotic treatment (3–5). Elevated inflammatory cytokines, microglial activation, and altered immune signaling pathways have been associated with synaptic dysfunction and impaired neural plasticity in SCZ (3). These findings have prompted growing interest in therapeutic strategies targeting neuroplasticity and immune-related mechanisms, including compounds with anti-inflammatory and neuromodulatory properties, as potential avenues for addressing treatment-refractory disease processes.

Astrocytes have emerged as central regulators of synaptic plasticity and neuroimmune signaling, extending far beyond their traditional role as passive support cells. As the most abundant glial cell type in the mammalian brain, they actively modulate glutamatergic neurotransmission, synapse formation and maintenance, neural metabolism, and inflammatory responses through the release and uptake of cytokines, chemokines, and neuromodulatory factors (6,7). Increasing evidence implicates astrocytic dysfunction in SCZ, including altered inflammatory gene expression, impaired glutamate homeostasis, and disrupted regulation of synaptic connectivity (8–11). Such abnormalities may contribute to excitatory-inhibitory imbalance and impaired neuroplasticity observed in affected individuals. However, mechanistic investigations into the cellular pathology of SCZ have been limited by restricted access to living human brain tissue and the translational limitations of animal models. Human induced pluripotent stem cell (iPSC)-based systems offer a powerful alternative, enabling patient-relevant modeling of neurodevelopmental and cellular processes (12). In particular, iPSC-derived human cortical spheroids (hCS) recapitulate key aspects of early cortical development and organization (13), while iPSC-astrocytes provide a tractable platform for investigating glial contributions to neuroinflammation, synaptic regulation, and disease-relevant molecular phenotypes in vitro (12,14).

Muscimol, the principal psychoactive compound of *Amanita muscaria*, is a potent and selective agonist of GABA_A_ receptors, structurally resembling the endogenous inhibitory neurotransmitter GABA (15,16). Unlike benzodiazepines and barbiturates, which act as allosteric modulators, muscimol binds directly to the orthosteric GABA_A_ receptor site, producing robust inhibitory effects on neuronal excitability (17). GABA_A_ receptors are widely expressed in cortical and hippocampal circuits and are also highly expressed in cortical astrocytes, where they regulate glutamatergic signaling, inflammatory responses, and synaptic homeostasis (18). Previous reports demonstrate that muscimol enhances cognition, memory and learning while reducing markers of neuroinflammation and astrocyte reactivity, including GFAP expression (19–21). Furthermore, GABA_A_ receptor activation by muscimol protects against neurotoxicity and restores network stability in experimental models of neurodegeneration (22,23). These findings suggest that direct GABAergic modulation may represent a promising strategy for restoring neuroplastic and neuroimmune balance in disorders characterized by excitatory-inhibitory and inflammatory dysregulation, such as SCZ (3,24–26). However, the functional and cell type-specific effects of muscimol on these pathways in SCZ remain unexplored.

In the present study, we aimed to map the acute and long-term effects of muscimol on iPSC-derived hCS and astrocytes on neuroplasticity and neuroinflammation, in order to identify potential new drug targets in SCZ. Using transcriptomic, proteomic, and functional analyses, we comprehensively examined the effects of muscimol on signaling pathways related to synaptic plasticity and neuroimmune regulation. To further resolve cell type-specific mechanisms, we integrated these findings with postmortem brain transcriptomic datasets and cell type deconvolution approaches. This strategy enabled identification of disease and cell type-specific factors through which GABA_A_ receptor activation modulates neuroplastic and inflammatory processes relevant to SCZ pathophysiology.

## MATERIALS AND METHODS

### Sample characteristics

Skin biopsy donors were recruited through the Norwegian TOP (Thematically Organized Psychosis) study. Recruitment procedures, inclusion and exclusion criteria, and clinical assessments for the TOP study have been described elsewhere (27,28). Fibroblasts were isolated from 10 donors with SCZ and 7 healthy controls (CTRL), selected based on clinical information and SCZ polygenic load. Cases and controls were matched on age and sex. Supplementary Table S1 summarizes the clinical characteristics of the study participants.

### IPSC derivation and characterization

Skin fibroblasts were cultured and reprogrammed as previously described (29). Each iPSC line was subjected to rigorous quality control by phenotyping, regular monitoring of morphology, and pluripotency marker expressions at the Norwegian Core Facility for Human Pluripotent Stem Cell Research Centre. KaryoStat GeneChip array (Thermo Fisher Scientific) was used for karyotyping of iPSCs at passages 15 to 20 for monitoring chromosomal aberrations.

### Generation of cortical spheroids

IPSCs differentiation to human cortical spheroids (hCSs) was done as published previously (30,31), following a protocol with proven high reproducibility (32,33). Briefly, iPSCs were dissociated into single cells with Accutase (Sigma-Aldrich) and added to a 6-well Aggrewell 800 (STEMCELL Technologies), centrifuged for 3 minutes at 100g, and incubated at 37 °C with 5% CO2 for 24 hours. Afterward, hCSs were collected and transferred into ultra-low-attachment plastic dishes (Thermo Fisher Scientific) in Essential 6 Medium (Thermo Fisher Scientific) supplemented with dual SMAD inhibition (SB-431542 10□μM; Tocris Bioscience, and dorsomorphin 2.5□μM, Sigma-Aldrich) and Wnt inhibitor XAV-939 (2.5□μM, Tocris Bioscience). Medium was changed daily from day 2 to day 6 (E6 plus inhibitors). After day 6, suspended spheroids were shifted to neural medium: Neurobasal A (Life Technologies, 10888), GlutaMax (1:100, Life Technologies, 35050) and B-27 supplement without vitamin A (Life Technologies, 12587). Neural medium was supplemented for 19 days until day 24 with 20□ng/ml EGF (R&D Systems, 236-EG) and 20□ng/ml bFGF (R&D Systems, 233-FB), with medium changed daily in the first 10 days and every other day for the following 9 days. From day 24 until day 43 neural medium was supplemented with 20□ng/ml BDNF (Peprotech, 450-02) and 20□ng/ml NT3 (Peprotech, 450-03) to promote neural differentiation, with medium changed every other day. From day 43 onwards, only neural medium without growth factors was changed every 3-4 days. The hCSs were differentiated until day 240 before starting treatment with muscimol and IL-1β.

### Differentiation of iPSC-astroglia

For the current work, we used a previously established, robust hiPSC-astrocyte monoculture model with high culture purity. Astrocytes were differentiated from n=4 SCZ and n=4 CTRL donors of the original iPSC pool above, following an optimized glial differentiation protocol (8,9), and characterized using rigorous phenotyping and functional assays as reported previously (34). Briefly, at passages 23-25, iPSC colonies were transferred to 6-well tissue culture plates in NMM medium consisting of 50 % DMEM/F12 and 50 % Neurobasal medium (both from Thermo Fisher, Waltham, MA, US) supplemented with 0.5 % (v/v) N2, 1 % (v/v) B27 (both from Invitrogen, Carlsbad, CA, US), 5 μg/ml human insulin, 40 ng/ml triiodothyronine (T3), 10 μM β-mercaptoethanol, 1.5 mM L-glutamine, 100 μM NEAA, 100 U/ml penicillin and 100 μg/ml streptomycin (all from Sigma-Aldrich, St. Louis, MO, USA). 20 ng/ml EGF and 4 ng/ml bFGF (both from Peprotech, Rocky Hill, NJ, USA) were used in the cultures to establish optimal growth and differentiation conditions. The astroglia induction phase included days 1-42 using the N2/B27 protocol. On day 42, NMM medium was switched to ScienCell AM medium (ScienCell, Carlsbad, CA, USA) for time of experiments. For phenotyping, the mRNA expression of *PAX6*, *Nestin*, and the astrocyte-specific markers *GFAP*, *S100B*, *SLC1A2*, *SLC1A3*, *FABP7*, *AQP4*, *ALDH1L1*, and *ALDOC* was monitored using a custom-made TLDA gene array card (Thermo Fisher) on a qPCR platform (8). For additional quality control, cells were stained with anti-GFAP, anti-S100B and anti-AQP4 antibodies (all from Abcam, Cambridge, UK) and were analyzed using fluorescent microscopy on day 42 (34). Functional characterization of differentiated astrocytes was performed as described previously (34). For the details of selected donors, see Suppl. Table S2.

### In vitro treatment paradigms

To assess the transcriptional effects of inflammation, day 240 hCSs were treated with 10 ng/ml human recombinant IL-1β (Peprotech, NJ, US) for 8h. To assess the effects of muscimol in in vitro cultures, cells were treated with 1 μM muscimol for 8h (transcriptional studies), 24h (secreted factors), or 7 days (chronic effects). Immune challenge of astrocytes was done using the synthetic viral dsRNA mimic polyadenylic-polyuridylic acid (poly(A:U)) at a working concentration of 500 ng/ml for 24h (InvivoGen, CA, US). For cytoplasmic delivery of poly(A:U), the LyoVec system was used following the manufacturer’s recommendation (InvivoGen). To ihibit GABA_A_ receptor sites, bicuculline (Tocris, Bristol, UK) was used at a working concentration of 10 μM, based on a previous protocol (35). Cell pellets and culture supernatants were collected after 8h of incubation for RNA-Seq, and after 24h of incubation for ELISA, MSD, and glutamate uptake analyses. To assess the chronic, long-term transcriptional effects of muscimol on hCSs, cell pellets were collected 7 days post-treatment. Harvested samples were either immediately used for analyses, or stored at −80 °C. For hCS experiments, in each case, one hCS per donor and treatment modality was used. To assess hCS volumes for the normalization of ELISA and MSD assays, images were taken and ImageJ software was used for area counting. Images were converted to 8-bits and binary color, scale was set using as reference distance 34.8 mm of diameter of the culture dish; organoids were analyzed counting the area, and volume in mm³ was estimated using the formula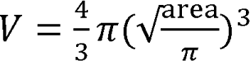

### RNA Extraction and Sequencing

Total RNA was extracted from all 85 hCS samples (17 donor iPSC lines, 5 treatment conditions each) using the RNeasy Plus Mini Kit (Qiagen). RNA yield was quantified with a NanoDrop 8000 Spectrophotometer (NanoDrop Technologies) and RNA integrity was assessed with Bioanalyzer 2100 RNA 6000 Nano Kit (Agilent Technologies). Library preparation and paired-end RNA-sequencing were carried out at the Norwegian High-Throughput Sequencing Centre (www.sequencing.uio.no). Briefly, libraries were prepared with the TruSeq Stranded mRNA kit from Illumina which involves poly-A purification to capture coding as well as several non-coding RNAs. The prepared samples were then sequenced in one batch on an Illumina NovaSeqX platform at an average depth of 70 million reads per sample, using a paired-end read length of 150 bp.

### Data Processing

Raw sequencing reads were quality assessed with FastQC (Babraham Institute). To pass the initial QC check, the average Phred score of each base position across all reads had to be at least 30. Reads were further processed by cutting individual low-quality bases and removing adapter and other Illumina-specific sequences with Trimmomatic (36) using default parameters. HISAT2 (36) was then used to first build a transcriptome index based on Ensembl annotations, and then to map the trimmed reads to the human GRCh38 reference genome. To quantify gene expression levels, mapped reads were summarized at the gene level using featureCounts guided by Ensembl annotations (37).

### Differential expression (DE) analysis and Gene Ontology (GO) enrichment tests

Before conducting the DE analyses, genes were pre-filtered using the filtreByExpr function in the edgeR package (38). In addition, only protein coding and long non-coding RNAs, the two most abundant RNA species, were included. DE analyses were performed using limma-voom (39) together with the duplicateCorrelation and voomWithQualityWeights functions to account for the paired design in the treatment comparisons and to reduce outlier effects. For the analysis between baseline (untreated) CTRL and SCZ samples, sex and age were included as covariates in the model. An FDR value of <0.1 was used as the significance threshold. GO enrichment tests of significant DE genes were conducted with the over-representation analysis tool clusterProfiler (40) using the enrichGO function, selecting BP (biological processes) as the ontology of interest. A GO term was considered significantly enriched if the FDR was <0.1.

### RNA-seq of post-mortem brain samples (CMC)

Bulk RNA-seq data of postmortem brain samples (prefrontal cortex) were obtained from the CommonMind Consortium (CMC) (41). A subset of 551 samples (SCZ = 257, CTRL = 294) was used, excluding samples from the NIMH Human Brain Collection Core (HBCC) which were processed differently than the samples from the three other collections. Prefiltering and DE analysis were carried out as described above, adjusting for age, sex, and postmortem interval.

### Cell type deconvolution

Computational estimation of cell type abundances (deconvolution) and cell type-specific gene expression were performed with InstaPrism (42), a reimplementation of a Bayesian approach to the estimation of cell type composition (43). A primary single-cell dataset of the developing human neocortex was used as reference, excluding cell populations of non-neural lineage (44).

### In vitro assaying of secreted factors

Secreted levels of IL-6, TNF, IP10, BDNF, NGF, and MMP7 in hCS culture supernatants were analyzed by using a U-PLEX Biomarker kit from Meso Scale Discovery (Meso Scale Diagnostics, LLC) using a QuickPlex SQ120 instrument according to instructions from the manufacturer.

For ELISA cytokine assays, 2.5 × 10^5^ astrocytes were cultured in 1 ml AM medium in 12-well plate. Culture supernatants were harvested 24 h after 500 ng/ml poly(A:U) and/or 1 μM muscimol treatment, and the concentrations of IL-6, TNF, and interferon-β were measured using human cytokine ELISA kits (all from Thermo Fisher) following the manufacturer’s recommendations. The precision of the ELISA kits was the following: Intra-Assay variation: CV < 10 %; Inter-Assay variation: CV < 12 % (CV% = SD/meanX100).

### LC-MS/MS proteomics analysis of hCS organoids

Protein concentrations of GABA receptors and pathway elements in hCS lysates were analyzed using liquid chromatography-mass spectrometry (LC-MS/MS) of non-treated hCSs. Cells were lysed in RIPA buffer (Sigma) with protease and phosphatase inhibitor cocktail (Thermo Scientific) and 5 mM EDTA (Thermo Scientific). Total protein concentration was measured with BCA protein assay kit (ThermoFisher). LC-MS/MS analysis was carried out using an EVOSEP one LC system (EVOSEP Biosystems, Denmark) coupled to a timsTOF Pro2 mass spectrometer, using a CaptiveSpray nano electrospray ion source (Bruker Corporation, Germany). 4 ul of digested peptides were loaded onto a capillary C18 column (15 cm length, 150 μm inner diameter, 1.5 μm particle size, EVOSEP, Odense Denmark). Peptides were separated at 40 °C using the standard 30 sample/day method from EVOSEP. The timsTOF Pro2 mass spectrometer was operated in DIA-PASEF mode. Mass spectra for MS were recorded between m/z 100 and 1700. Ion mobility resolution was set to 0.85-1.30 V·s/cm over a ramp time of 100 ms. The MS/MS mass range was limited to m/z 475-1000 and ion mobility resolution to 0.85-1.27 V s/cm to exclude singly changed ions. The estimated cycle time was 0.95 s with 8 cycles using DIA windows of 25 Da. Collisional energy was ramped from 20eV at 0.60 V s/cm to 59eV at 1.60 V s/cm. Raw data files from LC-MS/MS analyses were submitted to DIA-NN (version 1.8.1) for protein identification and label-free quantification using the library-free function. The UniProt human database (UniProt consortium, European Bioinformatics Institute, EMBL-EBI, UK) was used to generate library *in silico* from a human FASTA file. Carbamidomethyl(C) was set as a fixed modification. Trypsin without proline restriction enzyme option was used, with one allowed miscleavage and peptide length range was set to 7-30 amino acids. The mass accuracy was set to 15ppm, and precursor false discovery rate (FDR) allowed was 0.01 (1%). LC-MS/MS data quality evaluation and statistical analysis was done using software Perseus ver 1.6.15.0.

### Glutamate uptake analysis

Indirect assessment of glutamate uptake by iPSC-derived astrocytes was done using a colorimetric glutamate assay kit following the manufacturer’s recommendations (Abcam), as described in details elsewhere (34). Briefly, cells were incubated in AM medium at 37 °C and 5 % CO2 for 24 h then supernatants were collected and assayed. Native AM cell medium (medium control) and donor/clone-specific iPSC lines were used as controls.

### Statistics

All statistical analyses were conducted using Python (v3.x) with the statsmodels and SciPy libraries. For the organoid MSD datasets, analyses were tailored to account for repeated measurements obtained from the same donor-derived cell lines across multiple experimental conditions. Cytokine and growth factor concentrations were analyzed using linear mixed-effects models to appropriately account for within-donor correlations arising from repeated measurements. For each analyte, diagnosis (SCZ vs CTRL), condition, and their interaction were included as fixed effects, while donor was modeled as a random intercept. Planned pairwise comparisons versus the non-treated (NT) condition were performed to assess treatment effects within diagnostic groups. Where appropriate, Holm-adjusted p-values were calculated to control the family-wise error rate without overly penalizing statistical power. Estimated marginal means, model coefficients, 95% confidence intervals, and p-values were extracted for reporting. Data were visualized as mean ± SEM with individual donor values. For the ELISA (IL-6, TNF, IFNβ) and glutamate uptake measurements of astroglia cultures, population-averaged effects were estimated using generalized estimating equations (GEE) with a Gaussian distribution and exchangeable working correlation structure to account for repeated observations within donors. All tests were two-tailed, and statistical significance was defined as p < 0.05. Given the modest donor sample sizes typical of iPSC studies, emphasis was placed on model-based estimation, effect directionality, and confidence intervals rather than sole reliance on null hypothesis significance testing.

## RESULTS

### Derivation of hCSs

For the current study, we used previously differentiated and characterized hCSs (30) derived from patients with SCZ and matched controls (see Methods). The hCSs were cultured for 240 days and were matched on age and sex (Suppl. Table S1). Importantly, SCZ donors were selected based on polygenic risk score (PRS) for SCZ, and mean PRS was significantly higher in the SCZ group (SCZ mean PRS = 1.53, CTRL mean PRS = −0.73, p = 0.0179; Suppl. Table S1).

### Muscimol modulates inflammatory and neuroplasticity-associated transcriptional programs in hCSs

To characterize the transcriptional effects of inflammatory and GABAergic modulation, we performed RNA sequencing on hCSs derived from patients (SCZ; n = 10) and healthy controls (CTRL; n = 7) following acute (8 h) and chronic (7-day) exposure to IL-1β and muscimol. Across all samples, acute IL-1β exposure induced robust upregulation of inflammatory and interferon-stimulated genes, including *MX1*, *CXCL10*, and *MMP7*, with Gene Ontology (GO) enrichment confirming activation of innate immune and cytokine signaling pathways, including defense response to virus and interferon signaling (Figure 1). Co-treatment with muscimol did not abolish this response but reduced the magnitude of inflammatory gene induction and simultaneously engaged transcriptional programs associated with macroautophagy, cellular stress adaptation, and transcriptional regulation (Fig. 1B-C). In contrast to IL-1β, muscimol exposure alone induced transcriptional programs distinct from inflammatory activation. Acute muscimol (MA) treatment primarily affected regulatory and signaling-associated genes, whereas chronic muscimol (MC) treatment induced upregulation of the activity-dependent transcription factor *ELK1* and enrichment of biological processes associated with cytoskeletal organization, calcium handling, and notably, regulation of neuron projection and synaptic development, consistent with activation of neuroplasticity-associated transcriptional programs (Fig. 1B-C).

**Figure 1.**
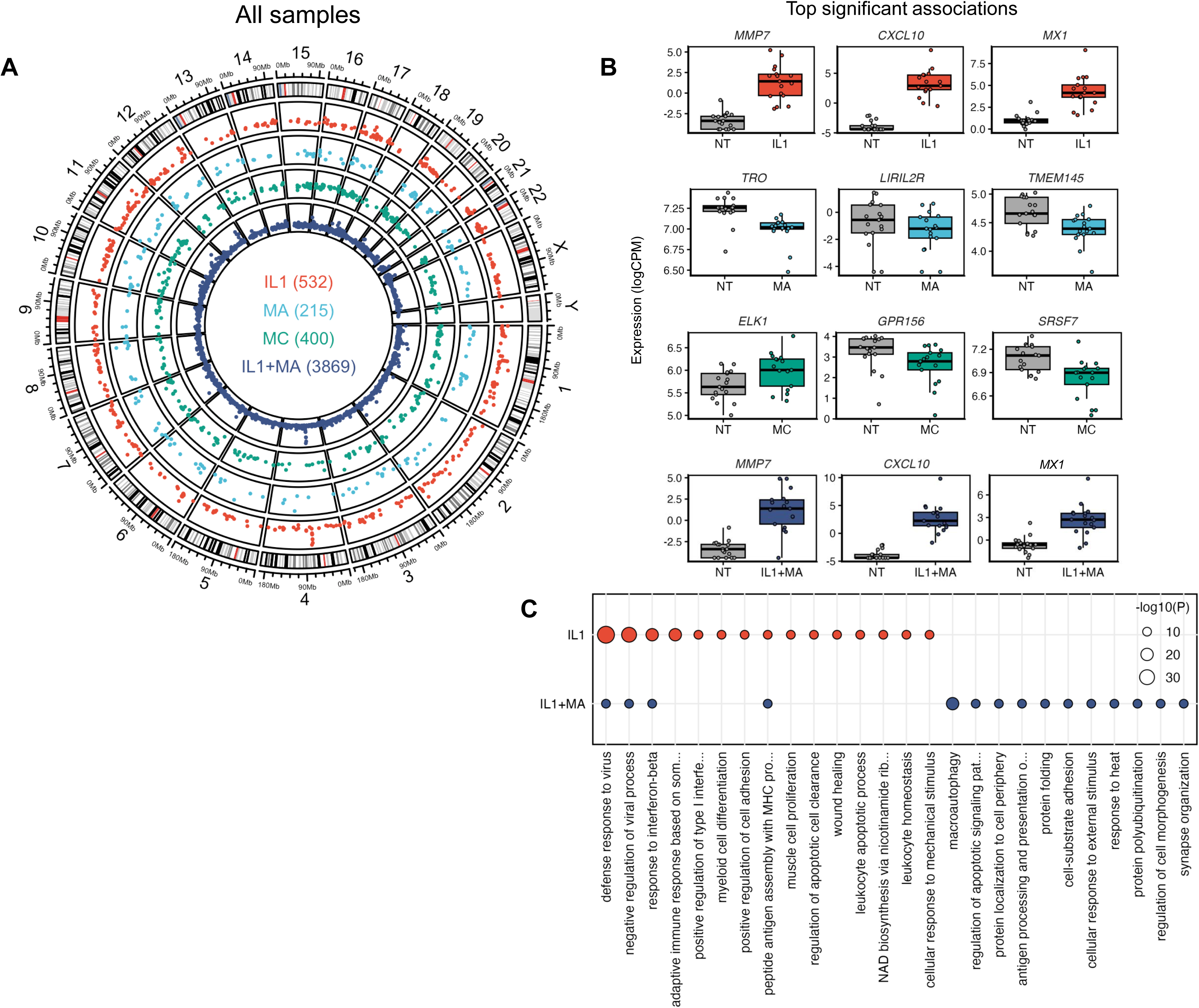
Transcriptional effects of muscimol and inflammatory modulation in hCSs. **(A)** Circos plot showing significant DE genes per chromosome per treatment for all samples cobined. **(B)** Top significant genes per treatment. **(C)** GO enrichment analysis of significant genes per treatment. No significant GO terms were identified for MA and MC. Treatment conditions: IL1: 10 ng/ml IL-1β, 8h; MA: muscimol/acute, 1 μM/ml muscimol, 8h; MC: muscimol/chronic, 1 μM/ml muscimol, 7 days; IL1+MA: co-treatment with IL-1β and muscimol, as above.

Stratified analyses revealed both shared and diagnosis-dependent responses. In CTRL-derived spheroids, muscimol induced transcriptional programs enriched for cytoskeletal remodeling, actomyosin organization, and intracellular calcium regulation, including upregulation of genes such as *TTN*, *SRL*, and *ATP2A1* (Fig. 2B-C), indicating coordinated structural and plasticity-associated transcriptional responses (45) (Figure 2). MC exposure further amplified enrichment of pathways associated with structural organization and neuron projection development (Fig. 2C). In SCZ-derived spheroids, IL-1β similarly induced inflammatory gene expression, including upregulation of *MX1*, *CXCL10*, and *ICAM1* (46) (Figure 3). Muscimol exposure in SCZ spheroids induced transcriptional changes involving regulatory and signaling-associated genes, including acute upregulation of *PLN*, a regulator of intracellular calcium homeostasis (47), and chronic upregulation of signaling-related genes including *FLT1*, alongside enrichment of regulatory pathways including ERK signaling and cellular stress response processes (Fig. 3B-C). As observed in pooled analyses, muscimol co-treatment attenuated IL-1β-induced transcriptional responses while engaging adaptive transcriptional programs.

**Figure 2.**
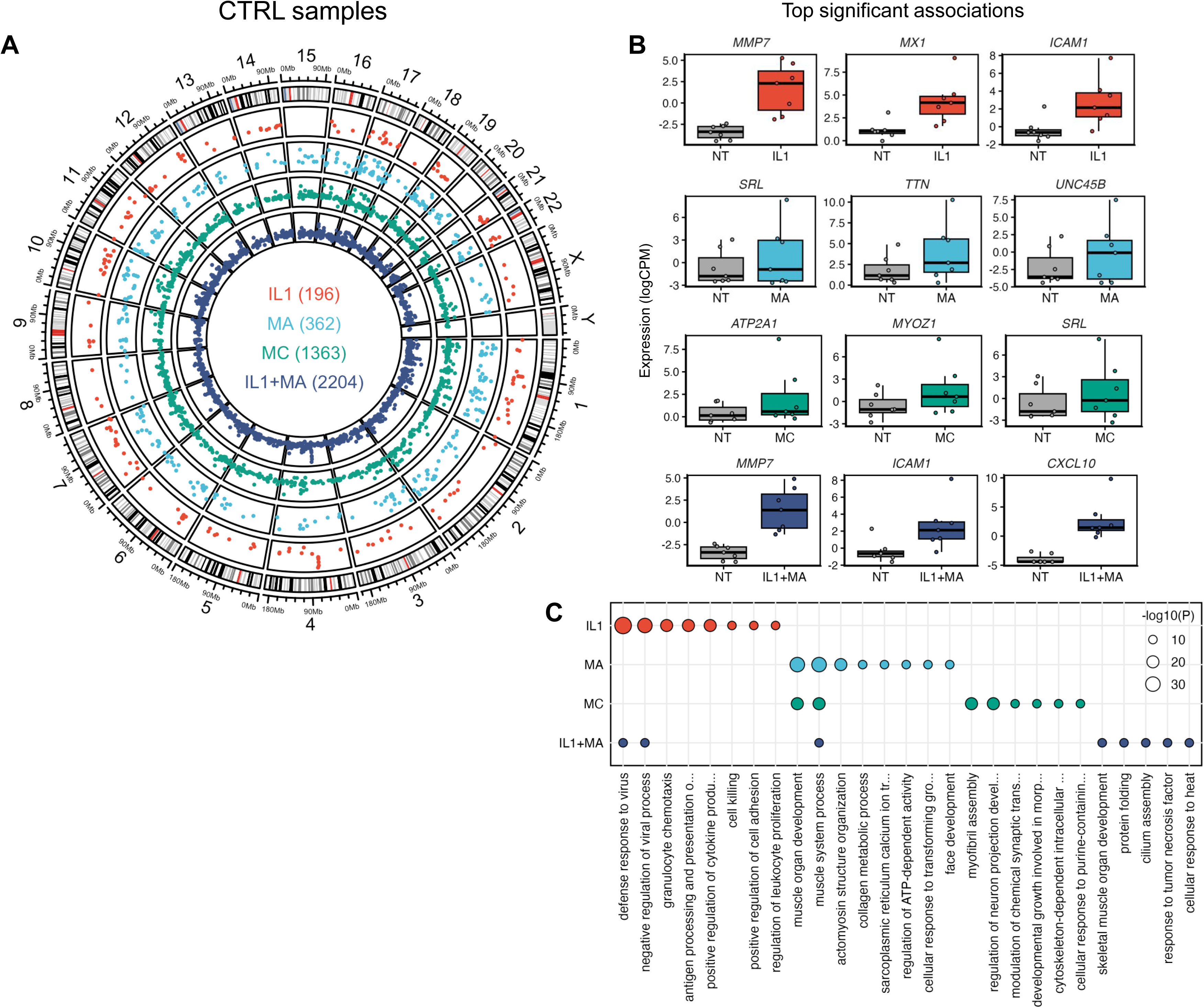
Transcriptional effects of muscimol and inflammatory modulation in CTRL hCSs. **(A)** Circos plot showing significant DE genes per chromosome per treatment in CTRL. **(B)** Top significant genes per treatment. **(C)** GO enrichment analysis of significant genes per treatment. Treatment conditions: IL1: 10 ng/ml IL-1β, 8h; MA: muscimol/acute, 1 μM/ml muscimol, 8h; MC: muscimol/chronic, 1 μM/ml muscimol, 7 days; IL1+MA: co-treatment with IL-1β and muscimol, as above.

**Figure 3.**
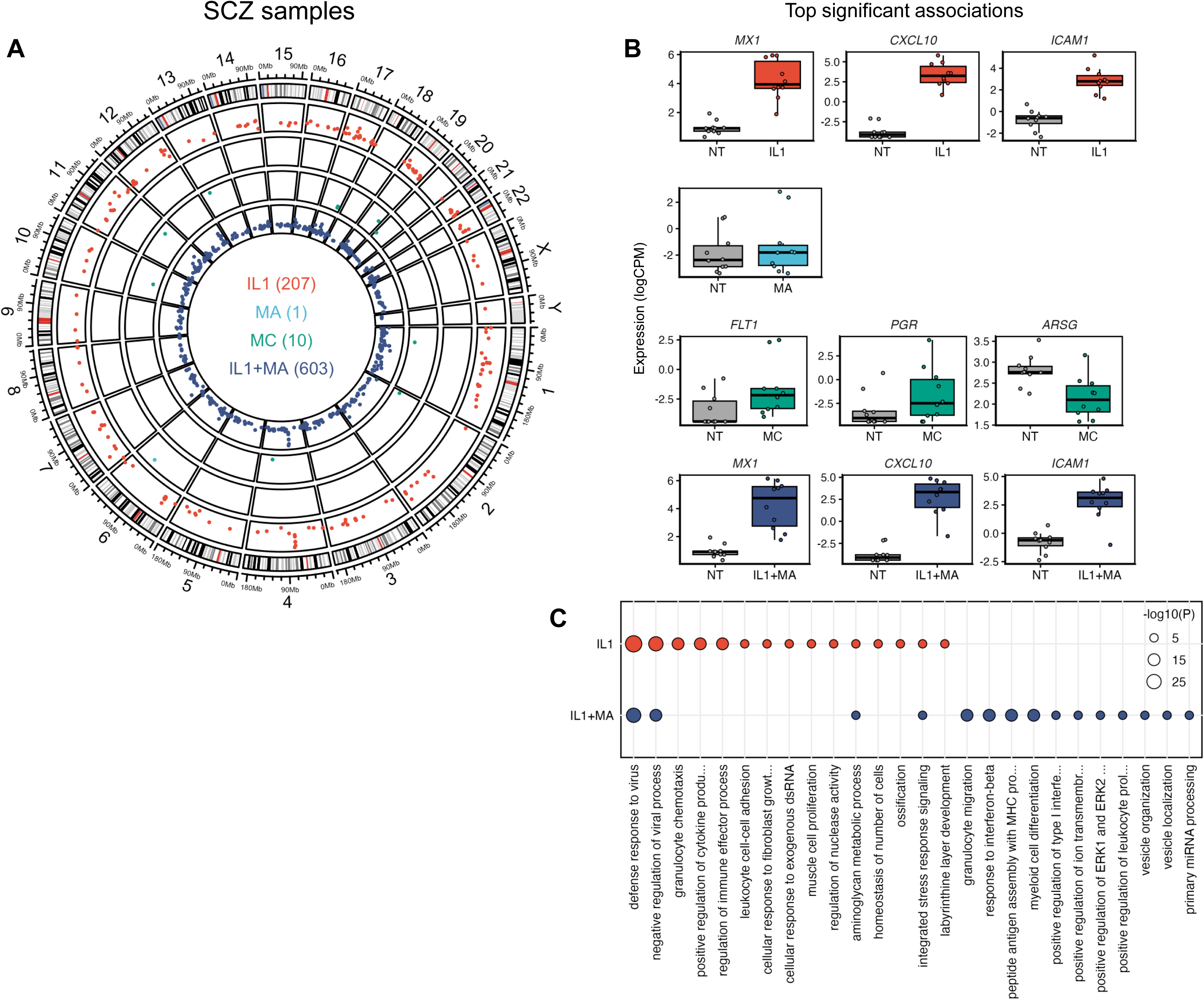
Transcriptional effects of muscimol and inflammatory modulation in SCZ hCSs. **(A)** Circos plot showing significant DE genes per chromosome per treatment in SCZ. **(B)** Top significant genes per treatment. **(C)** GO enrichment analysis of significant genes per treatment. No significant GO terms were identified for MA and MC. Treatment conditions: IL1: 10 ng/ml IL-1β, 8h; MA: muscimol/acute, 1 μM/ml muscimol, 8h; MC: muscimol/chronic, 1 μM/ml muscimol, 7 days; IL1+MA: co-treatment with IL-1β and muscimol, as above.

To further investigate whether muscimol modulates expression of genes directly involved in GABAergic signaling and neuroplasticity, we performed targeted analysis of GABA receptor subunits, GABA synthesis enzymes, and neurotrophin signaling components (Figure 4). MA exposure did not significantly alter expression of these genes in SCZ spheroids. However, MC treatment induced significant upregulation of *NTRK2* encoding TrkB, the neurotrophin receptor for BDNF, in SCZ spheroids (48) (Fig. 4A). In CTRL spheroids, MA treatment induced downregulation of GABA_A_ receptor subunits *GABRA5* and *GABRG3*, while chronic exposure further downregulated *GABRA5*, *GABRB1*, and *GABRG3*, as well as the neurotrophin gene *NGF* (Fig. 4A-B), indicating transcriptional regulation of inhibitory receptor composition and neurotrophin signaling following sustained GABA_A_ receptor activation (49).

**Figure 4.**
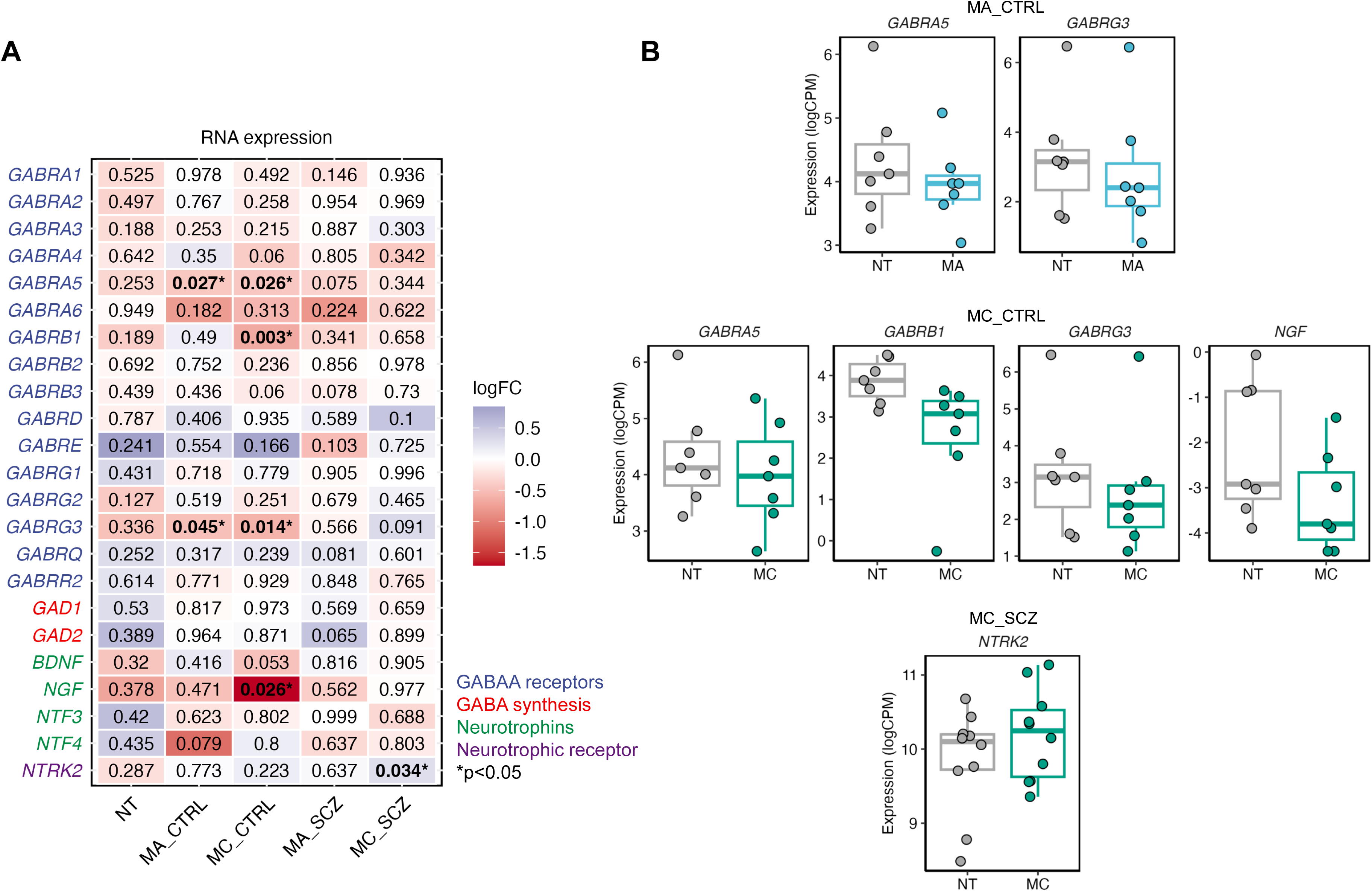
Transcriptional effects of muscimol treatment on neurotrophin and GABA signaling-related genes in hCSs. **(A)** Heatmap showing effect sizes (logFC) for comparisons between non-treated SCZ and CTRL samples (NT) and the different treatments for genes related to GABA and neurotrophin signaling. (B) Box plots of differentially expressed GABA and neurotrophin signaling. *p<0.05; NT: Non-treated (baseline) SCZ vs. CTRL; MA: muscimol acute (8h), MC: muscimol chronic (7 days).

Taken together, these results demonstrate that GABA_A_ receptor activation via muscimol induces transcriptional programs associated with neuroplasticity, structural remodeling and cellular adaptation, while simultaneously modulating inflammatory signaling. These effects involve both global transcriptional remodeling and targeted regulation of genes directly involved in inhibitory neurotransmission and neurotrophin-mediated plasticity.

### Baseline proteomic and human brain transcriptomic analyses reveal altered GABAergic and neurotrophin signaling in SCZ

To determine whether the transcriptional alterations in GABAergic and neuroplasticity-associated genes identified in hCSs were reflected at the protein level and in human brain tissue, we performed targeted proteomic analysis of cortical spheroids and examined RNA expression of GABAergic and neurotrophin signaling genes in dorsolateral prefrontal cortex (DLPFC) samples from the CMC.

Using LC-MS analysis of total protein lysates from CTRL and SCZ hCSs, we quantified baseline protein expression of key GABA receptor subunits, GABA synthesis enzymes, and neurotrophin receptors. Among measurable targets, the neurotrophin receptor NTRK2 exhibited significantly higher protein expression in CTRL compared to SCZ spheroids (FDR=0.014; Fig. 5). Additional GABAergic signaling components showed similar directional trends but did not reach statistical significance, including higher expression of GABRA3 and GABRG2 in CTRL hCSs and higher expression of GABRB3 and the GABA synthesis enzyme GAD1 in SCZ hCSs (Fig. 5). These findings indicate baseline alterations in neurotrophin receptor abundance and suggest broader differences in GABAergic signaling protein composition between CTRL and SCZ spheroids.

**Figure 5.**
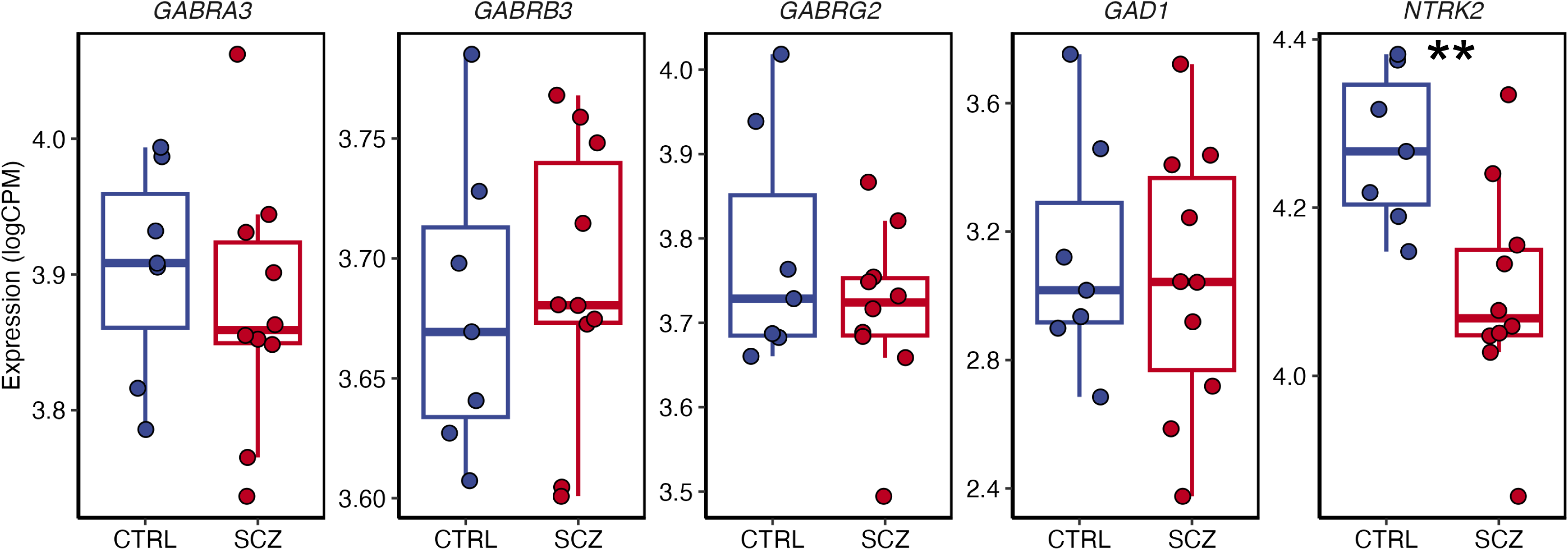
Proteomic characterization of GABA and neurotrophin signaling-related factors in hCSs. Box plots displaying LC-MS data of baseline protein expression in hCS, differentiated from healthy individuals (CTRL; n=7) or from patients with schizophrenia (SCZ; n=10). **FDR=0.014.

To assess whether similar alterations were present in human brain tissue, we analyzed RNA expression of GABAergic and neurotrophin signaling genes in postmortem DLPFC samples from the CMC (CTRL, n=294; SCZ, n=257). SCZ samples exhibited modest but significant differential expression of multiple GABA_A_ receptor subunits, including increased expression of *GABRA2*, *GABRA5*, *GABRB1*, *GABRB3*, and *GABRQ*, and decreased expression of *GABRD* (FDR<0.05; Fig. 6). Additional receptor subunits, including *GABRA3*, *GABRA4*, and *GABRG3*, also showed significantly increased expression in SCZ samples (p<0.05; Fig. 6), indicating widespread alterations in inhibitory receptor expression in SCZ cortex.

**Figure 6.**
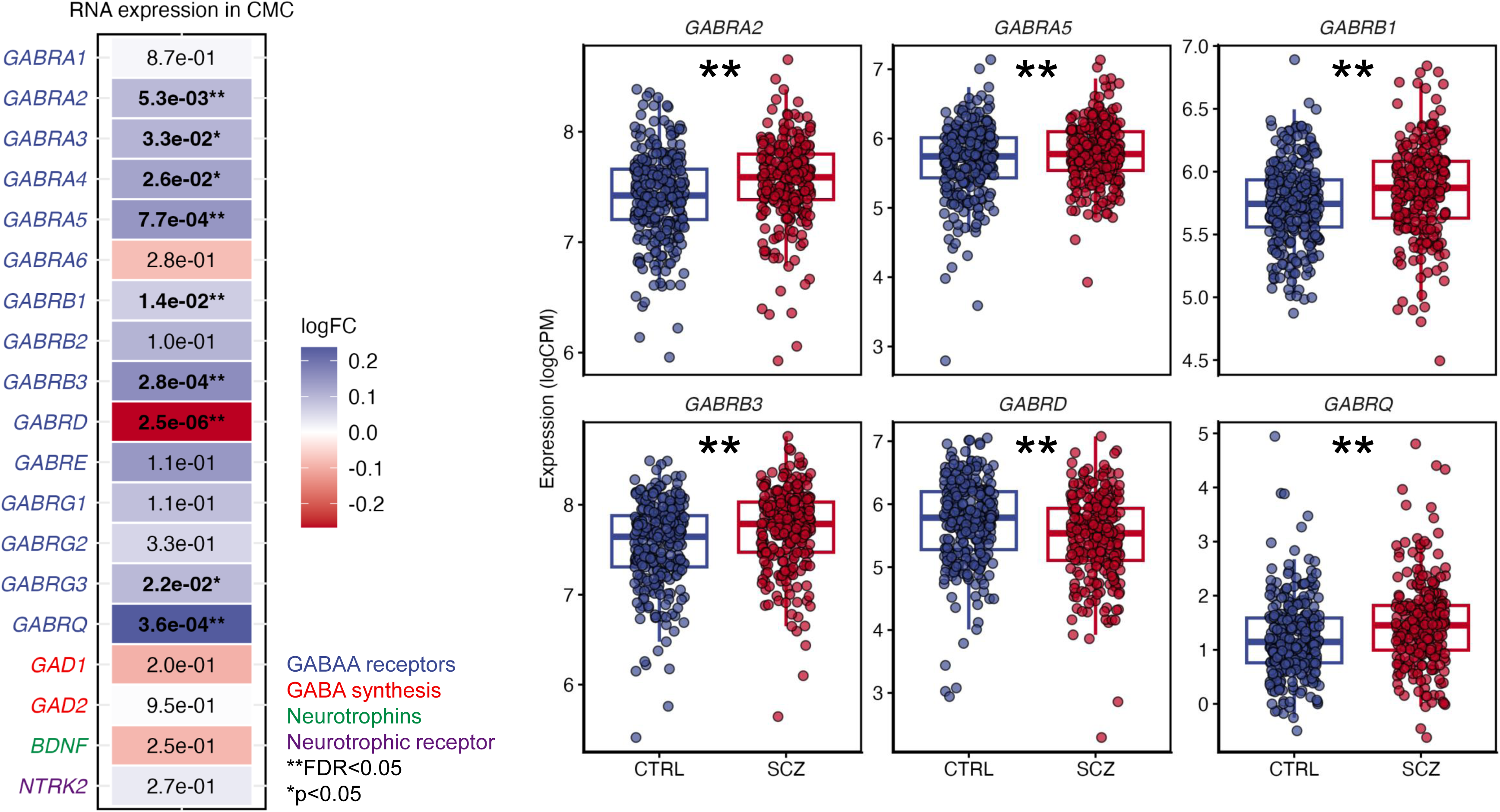
Gene expression of GABA and neurotrophin-signaling related genes in human postmortem brain samples. Left panel: Expression of genes related to GABA and neurotrophin signaling in human prefrontal cortex samples from the CMC (CTRL, n=294; SCZ, n=257). **Right panel:** Visualization of the top six significantly associated DE denes extracted from the left panel. Baseline mRNA expression of factors in brains in SCZ vs. CTRL; **FDR<0.05, *p<0.05.

Together with the transcriptomic findings in hCSs (Figs. 1-4), these results suggest alterations in GABAergic and neurotrophin signaling pathways across transcriptional and proteomic levels. In particular, the reduced baseline NTRK2 protein expression observed in SCZ hCSs contrasts with the transcriptional induction of *NTRK2* following chronic muscimol exposure (Fig. 4), suggesting altered neurotrophin signaling regulation and adaptive capacity. Furthermore, the widespread differential expression of GABA receptor subunits observed in human DLPFC samples supports the presence of altered inhibitory signaling architecture in SCZ. Collectively, these findings indicate that both neurotrophin signaling and GABAergic receptor expression are altered in SCZ and are dynamically regulated by GABA_A_ receptor activation, supporting a role for impaired inhibitory and neuroplasticity-associated signaling in the transcriptional and proteomic phenotype observed in SCZ hCSs.

### Muscimol modulates inflammatory and neurotrophic signaling in hCSs

To determine whether muscimol-induced transcriptional programs translated into functional protein-level changes, we quantified secretion of inflammatory cytokines (IL-6, TNF, IP-10), neurotrophins (BDNF, NGF), and the extracellular matrix remodeling enzyme MMP-7 in organoid supernatants under baseline, inflammatory, and muscimol treatment conditions (Fig. 7).

**Figure 7.**
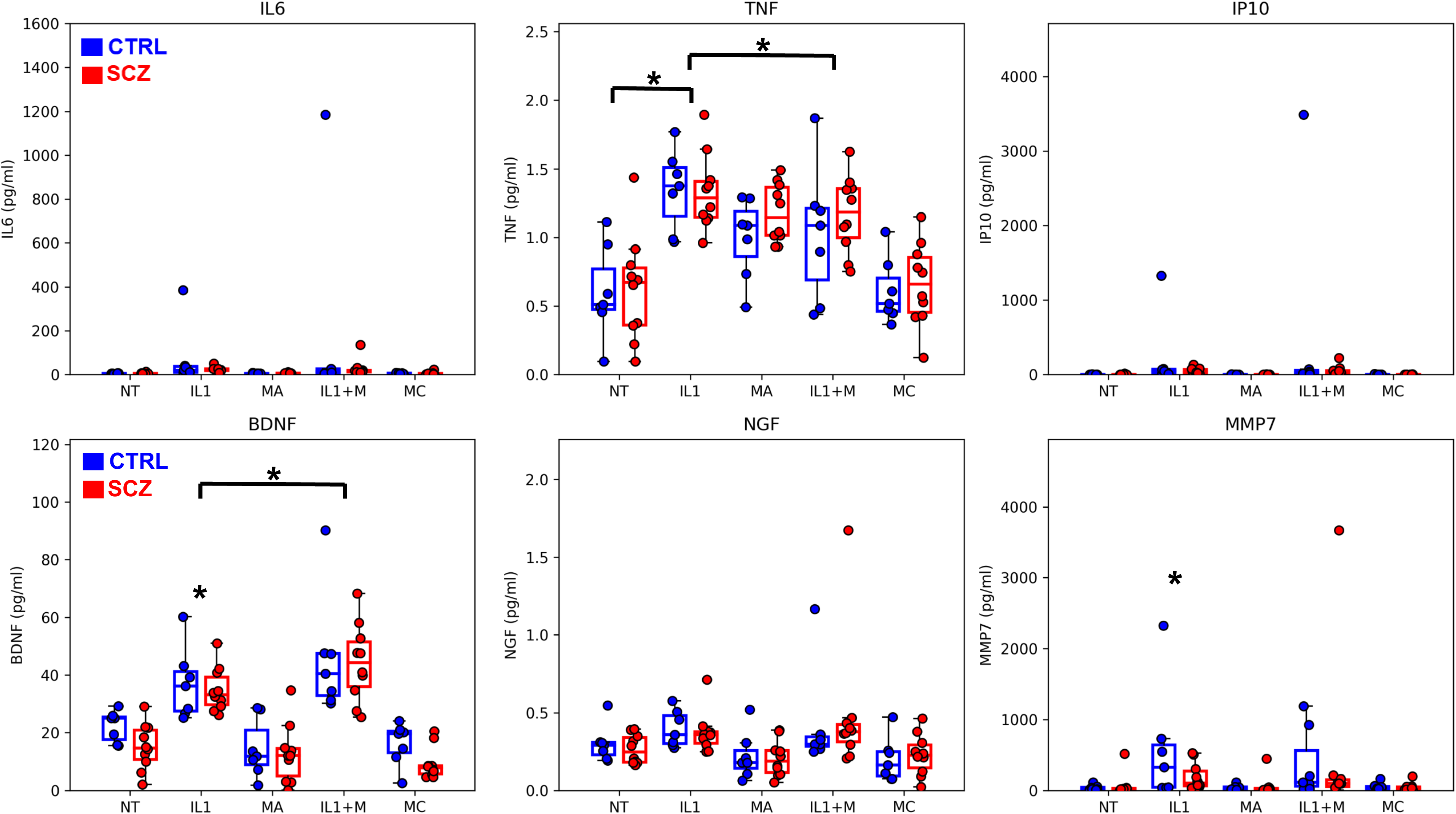
Biomarker profile of hCS supernatants following muscimol treatment and immune modulation. Levels of secreted factors in human hCS cultures following inflammatory modulation and/or muscimol treatment. Inflammatory cytokines (IL-6, TNF, IP10), neurotrophins (BDNF, NGF), and factors related to extracellular matrix remodeling (MMP7) were assayed using MSD (for details, please see Methods). NT: non-treated control; IL1: 10 ng/ml IL-1β, 24h; MA: muscimol/acute, 1 μM/ml muscimol, 24h; IL1+M: co-treatment with 10 ng/ml IL-1β and 1 μM/ml muscimol muscimol, 24h; MC: muscimol/chronic, 1 μM/ml muscimol, 7 days. *p<0.05, ***p<0.005; for details on statistical analyses, please see Methods, and Suppl. Table S3.

Under baseline conditions, MA treatment did not increase secretion of inflammatory cytokines compared to non-treated (NT) cultures. IL-6, TNF, and IP-10 remained low and comparable to NT in both CTRL and SCZ organoids, indicating that acute GABA_A_ receptor activation alone does not induce inflammatory signaling (Fig. 7). MC treatment similarly did not elevate inflammatory cytokine secretion relative to NT, confirming that sustained GABA_A_ receptor activation does not trigger inflammatory activation at the organoid level (Fig. 7). In contrast, inflammatory stimulation with IL-1β induced the secretion of TNF, consistent with activation of innate immune-associated transcriptional programs identified in RNA-seq analyses (Figs. 1-3). Co-treatment with muscimol (IL-1+M) attenuated the secretion of TNF relative to the IL-1β-only condition. Because TNF expression is induced downstream of type I interferon signaling, which directly regulates expression of interferon-stimulated genes including *MX1*, these findings provide functional evidence that GABA_A_ receptor activation suppresses interferon-dependent inflammatory signaling pathways identified in transcriptomic analyses (Figs. 1-4).

Muscimol also modulated neurotrophin secretion. Co-treatment of IL-1β with muscimol increased BDNF production, consistent with parallel transcriptional regulation of neuroplasticity-associated signaling pathways under inflammatory conditions, including modulation of neurotrophin receptor expression, observed in transcriptomic analyses (Fig. 4). Interestingly, while IL-1β increased secretion of the extracellular matrix remodeling enzyme MMP-7, consistent with transcriptional induction of *MMP7*, muscimol co-treatment did not modulate MMP-7 secretion relative to IL-1β alone. In addition, no significant associations were observed accross diagnostic categories in any of the measured analytes (Fig. 7).

Together, these findings demonstrate that GABA_A_ receptor activation does not induce inflammatory signaling under baseline conditions but suppresses inflammatory and interferon-associated signaling following immune activation while simultaneously modulating neurotrophin pathways. These results provide functional validation of the interferon-associated transcriptional programs identified in RNA-seq analyses, including regulation of *MX1* and related interferon-responsive genes.

### Astrocytes mediate interferon-dependent immune and glutamatergic functional responses to muscimol

Computational deconvolution analysis of bulk RNA-seq data revealed that glutamatergic neurons and astrocytes represented the largest cellular fractions in 3D organoid cultures, with no significant differences in cell type proportions across treatment conditions (Fig. 8). These findings indicate that the observed transcriptional changes were not driven by shifts in cell type composition but instead reflect treatment-induced transcriptional regulation within resident cell populations.

**Figure 8.**
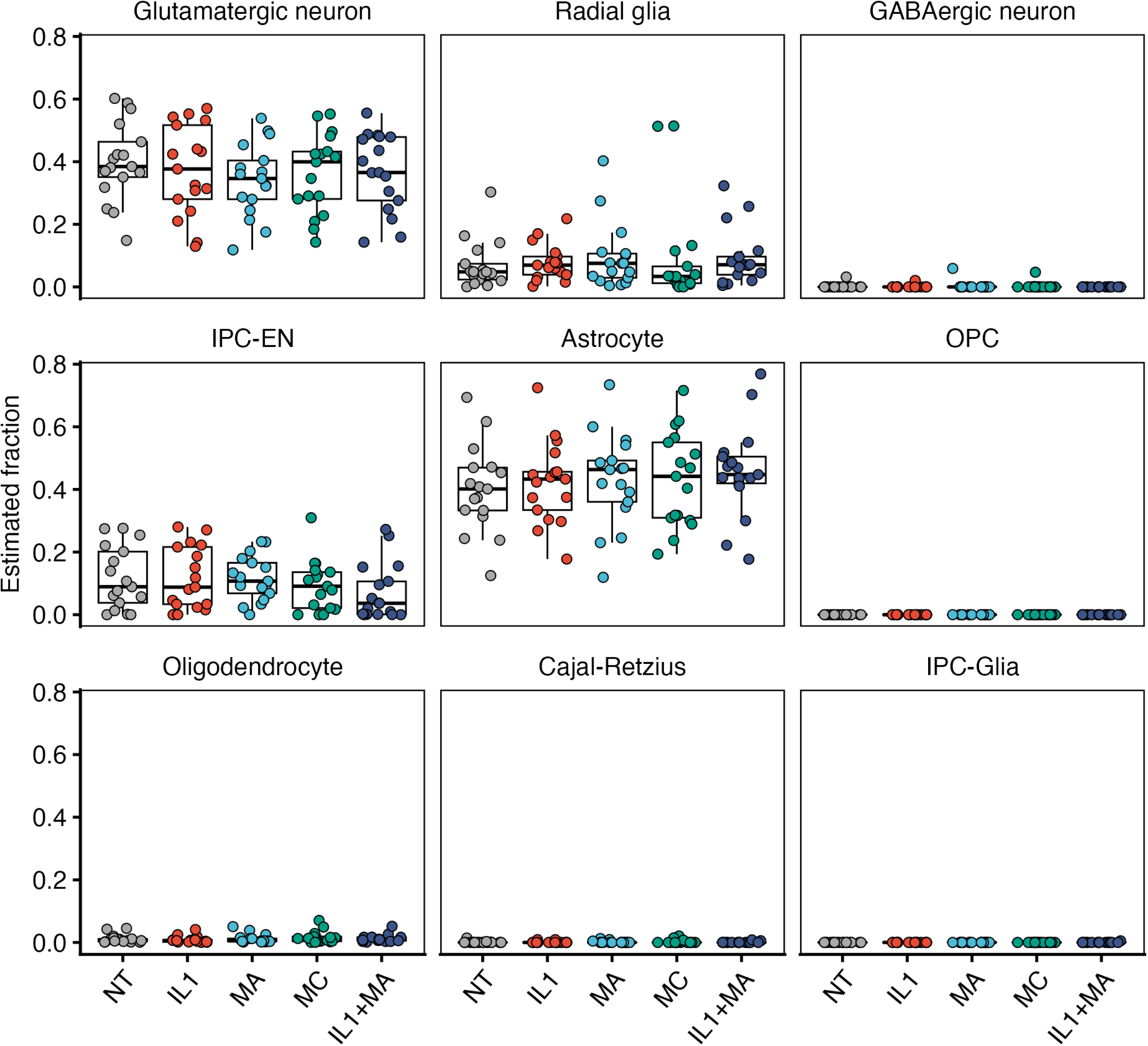
Cell type proportions in hCSs estimated by computational deconvolution. Glutamatergic neurons and anstrocytes were the cell populations displaying the largest fractions, but other cell types are also present in the cultures. No significant differences (t-test, p<0.05) were found in cell type fractions between NT and any of the treatment groups (in all hCS cultures). IPC-EN: intermediate progenitors of excitatory neurons; OPC: oligodendrocytes precursor cells.

Cell type-specific expression analysis further demonstrated that several of the most strongly regulated genes identified in bulk transcriptomic analyses exhibited cell type-specific expression patterns (Fig. 9). Notably, the interferon-stimulated gene *MX1*, one of the most significantly induced genes following inflammatory stimulation, showed prominent expression in astrocyte and glial populations (Fig. 9). Given that *MX1* expression is primarily regulated by type I interferons (50), these findings implicate astrocytes as key mediators of interferon-dependent transcriptional responses in hCSs.

**Figure 9.**
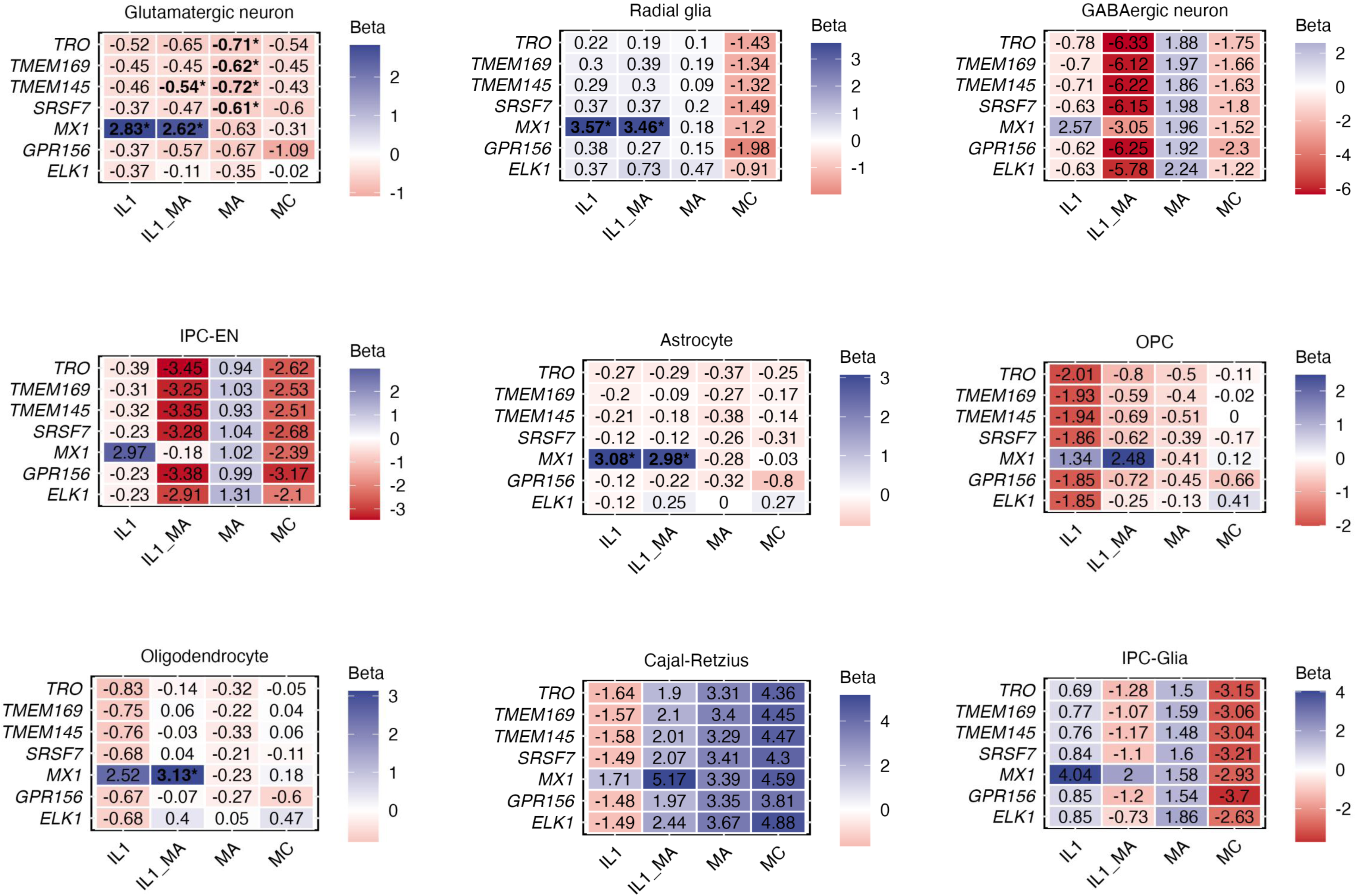
Cell type-specific expression analysis of the top 7 DE genes in hCSs. Multiple linear regression analysis adjusted for sex and age. *p<0.05/7 (Bonferroni correction for 7 analyzed genes). Beta values indicate expression levels in each treatment versus NT. Treatment conditions: IL1: 10 ng/ml IL-1β, 8h; MA: muscimol/acute, 1 μM/ml muscimol, 8h; MC: muscimol/chronic, 1 μM/ml muscimol, 7 days; IL1_MA: co-treatment with 10 ng/ml IL-1β and 1 μM/ml muscimol for 8h.

To directly assess astrocyte-specific functional responses, we generated iPSC-derived astrocytes from matched CTRL (n=4) and SCZ (n=4) individuals of the same donor pool, and performed immune and functional assays following viral mimic stimulation and muscimol treatment (Fig. 10). Poly(A:U) stimulation induced robust secretion of inflammatory cytokines, including IL-6, TNF, and particularly IFN-β, the principal upstream regulator of *MX1* expression (Fig. 10A-C). TNF and IFN-β secretion were significantly elevated in SCZ astrocytes compared to CTRL, indicating enhanced interferon-dependent immune activation in SCZ (Fig. 10C). Muscimol treatment attenuated inflammatory cytokine (TNF and IFN-β) secretion, demonstrating functional suppression of interferon-dependent immune signaling downstream of GABA_A_ receptor activation (Fig. 10B-C).

**Figure 10.**
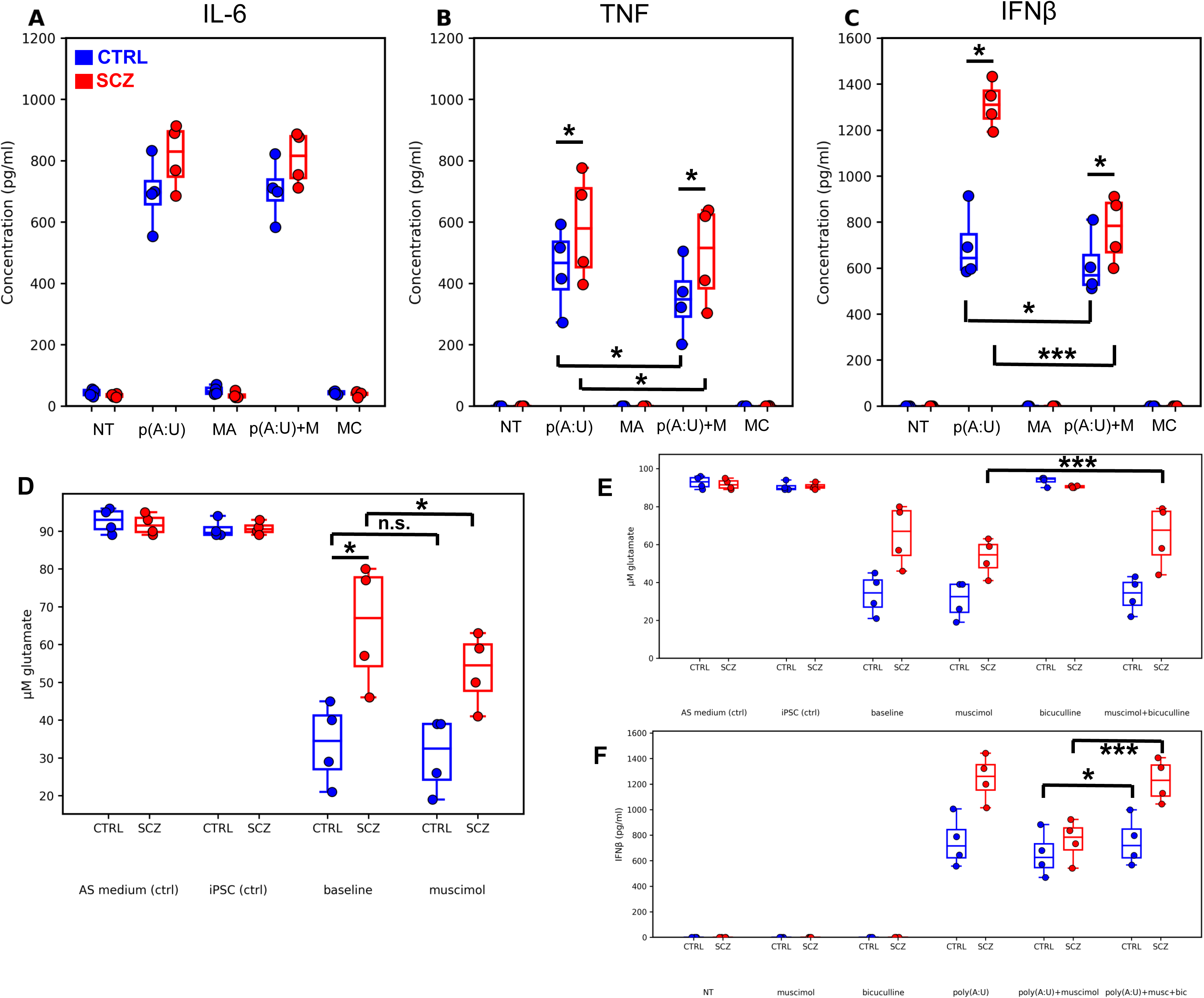
Functional and immune profiling of iPSC-astroglia following viral challenge and muscimol treatment. **(A-C)** ELISA assaying of inflammatory cytokines in iPSC-astrocyte cultures differentiated from healthy individuals (CTRL, n=4) or patients with schizophrenia (SCZ, n=4). **(D)** Glutamate uptake of resting/baseline and muscimol treated (24h) iPSC-astrocytes was assessed by a glutamate assay kit using the same donor lines as above (A-C). Native AM cell medium (AS medium ctrl), and donor/clone-specific iPSC lines were used as controls (iPSC ctrl). **(E-F)** Effects of bicuculline on glutamate uptake and IFNβ release. All assays and conditions are as detailed above and in Methods. Please note, that data for the control and muscimol values in Fig.10E are the same as those shown in Fig.10D and are included for reference to facilitate comparisons with the treated conditions (bicuculline and muscimol + bicuculline). NT: non-treated control; p(A:U): 500 ng/ml poly(A:U), 24h; MA: muscimol/acute, 1 μM/ml muscimol, 24h; p(A:U)+M: co-treatment with 500 ng/ml poly(A:U) and 1 μM/ml muscimol muscimol, 24h; MC: muscimol/chronic, 1 μM/ml muscimol, 7 days; bicuculline: 10 μM, in each case, treatment time adjusted to muscimol. *p<0.05, ***p<0.005, n.s.=non-significant; for details on statistical analyses, please see Methods, and Suppl. Tables S3-6.

In addition to immune effects, astrocytes exhibited altered glutamatergic functional phenotypes. SCZ astrocytes showed reduced glutamate uptake at baseline compared to CTRL astrocytes, indicating impaired glutamate handling (Fig. 10D). Muscimol treatment increased glutamate uptake in SCZ astrocytes, partly normalizing the observed deficit (Fig. 10D). These effects were blocked by the GABA_A_ receptor antagonist bicuculline, confirming that both anti-inflammatory and glutamate-regulatory effects were mediated through GABA_A_ receptor signaling in iPSC-astroglia (Fig. 10E-F).

Together, these findings identify astrocytes as key cellular mediators of interferon-dependent immune activation and glutamate dysregulation in SCZ-derived neural tissue, and demonstrate that muscimol, via GABA_A_ receptor activation, modulates astrocyte immune and functional phenotypes. These results provide functional validation of the interferon-associated transcriptional programs, including *MX1* induction, and neuroplasticity-associated signaling pathways identified in organoid transcriptomic and proteomic analyses.

## DISCUSSION

Schizophrenia is increasingly recognized as a disorder emerging at the intersection of neurodevelopment, immune dysregulation, and synaptic dysfunction. Converging epidemiological, genetic, and experimental evidence implicates inflammatory signaling, antiviral and interferon-mediated pathways in disease pathogenesis in a subset of patients (3,4,11). Prenatal exposure to viral infection, maternal immune activation, and early-life inflammatory insults significantly increase SCZ risk, supporting a neurodevelopmental model in which immune-mediated perturbations disrupt circuit maturation and synaptic stability (51–53). These processes involve glial cell activation, cytokine-mediated modulation of neurotransmission, and alterations in synaptic plasticity, thereby establishing long-term vulnerability to neuropsychiatric dysfunction.

Consistent with this framework, we observed robust activation of antiviral and interferon-responsive transcriptional programs in human cortical spheroids following inflammatory stimulation, prominently involving *MX1*, *CXCL10*, and *ICAM1*. *MX1*, a canonical interferon-stimulated gene, emerged as a central node across transcriptomic, cell-type-specific, and astrocyte functional analyses (50). This finding is notable because type I interferon signaling is a primary molecular response to viral infection and has been directly implicated in SCZ risk and pathophysiology (3,50). Astrocytes, which exhibited strong interferon-dependent responses and functional alterations, are key regulators of neuroimmune communication and synaptic homeostasis, suggesting that astrocytic antiviral signaling may represent a mechanistic link between environmental immune insults and persistent circuit dysfunction (8,10,11).

Activation of GABA_A_ receptors by the non-classic psychedelic muscimol exerted immunomodulatory and neuroplastic effects across our experimental systems. Muscimol attenuated inflammatory gene expression, reduced secretion of proinflammatory mediators, and normalized interferon-associated astrocyte responses, including IFNβ release. These findings align with prior evidence demonstrating that psychedelics modulate immune signaling through interactions with intracellular pathways regulating NF-κB, MAPK, and interferon signaling cascades, thereby controlling cytokine production and inflammatory gene expression (54,55). Such immunomodulatory effects are increasingly recognized as central components of psychedelic pharmacology and may contribute directly to therapeutic efficacy in various neurological and psychiatric disorders (56), including SCZ (57). In parallel, muscimol-induced transcriptional and functional signatures are also consistent with enhanced neuroplasticity. Chronic muscimol exposure upregulated *NTRK2* and *ELK1* while modulating genes associated with neuronal projection development and synaptic remodeling. Importantly, ELK1 has recently been implicated in psychiatric disorders, such as major depression, as a key factor promoting synaptic plasticity and maintaining neuroprotective responses against neurodegeneration (58,59). Understanding the etiology of SCZ remains a major challenge, with impaired neurotrophic support proposed to contribute to maladaptive neuronal network formation and reduced neuroplasticity (60). BDNF is a key regulator of neuronal survival and plasticity in major neurotransmitter systems, and its effects are primarily mediated through its high-affinity receptor TrkB/NTRK2, encoded by *NTRK2* (48). Disruptions in BDNF-NTRK2 signaling may therefore represent a central mechanism underlying structural and functional brain abnormalities in SCZ (60). Based on our results, the molecular pathways and factors underlying these neuroprotective and neuroplasticity-promoting effects appear to be directly modulated by muscimol. These findings therefore support the classification of psychedelics, including non-classic ones, as *psychoplastogens*, a class of compounds capable of rapidly promoting structural and functional neural plasticity (61,62). Psychoplastogens induce synaptic remodeling, increase dendritic spine density, and restore impaired neural circuitry, providing a mechanistic basis for sustained therapeutic effects. Consistent with this concept, muscimol also restored glutamate uptake deficits in SCZ-derived astrocytes, indicating functional rescue of impaired homeostatic processes critical for synaptic regulation (7).

Importantly, these effects occurred in the context of altered baseline GABAergic and neurotrophin signaling in SCZ-derived cortical tissue. Targeted transcriptomic and proteomic analyses revealed dysregulation of GABA receptor subunits and reduced NTRK2 protein expression in SCZ organoids, consistent with human postmortem data showing widespread alterations in inhibitory receptor expression (63). Independent validation in human dorsolateral prefrontal cortex samples demonstrated significant dysregulation of multiple GABA receptor subunits, confirming that inhibitory signaling abnormalities represent an important feature of SCZ pathophysiology consistent with previous reports and hypotheses (64–66). Our present findings suggest that impaired GABAergic signaling may contribute both to immune dysregulation and reduced plasticity capacity in SCZ.

The convergence of immune modulation and plasticity induction by muscimol provides a mechanistic framework linking neuroimmune dysfunction and synaptic pathology. Psychedelics have been shown to stimulate neurotrophic pathways, promote synaptic remodeling, and enhance neural circuit resilience (21,61). These neuroplastic and immunomodulatory effects likely reflect synchronized regulation of intracellular signaling pathways controlling synaptic stability, immune regulation, and cellular homeostasis. Notably, the ability of muscimol to suppress inflammatory interferon signaling while simultaneously restoring astrocyte function suggests that psychoplastogenic and immunomodulatory mechanisms are functionally intertwined. Our finding that muscimol potently suppresses type I interferon (IFN-β) responses in human iPSC-astrocytes identifies a previously unrecognized mechanism by which GABA_A_ receptor activation may directly regulate astrocyte-intrinsic innate immune signaling relevant to SCZ. Type I interferons have emerged as critical upstream amplifiers of neuroinflammation, capable of driving microglial activation, inducing complement cascade components, and promoting excessive synaptic pruning and neuronal element engulfment (67–69). Such interferon-dependent microglial phenotypes are highly phagocytic and have been implicated in maladaptive circuit refinement and synaptic loss, processes strongly linked to the neurodevelopmental and cognitive deficits characteristic of SCZ (70). Astrocytes represent a major source and regulator of type I interferons in the central nervous system and play a central role in orchestrating microglial activation states and neuroimmune homeostasis through cytokine-mediated cross-talk. Within this framework, the observed inhibitory effect of muscimol on astrocytic IFN-β production suggests that GABAergic signaling can directly constrain astrocyte-derived interferon tone, thereby limiting pathological astrocyte-microglia feed-forward inflammatory loops. This mechanism is particularly relevant given the growing recognition that astrocyte-mediated immune dysregulation may contribute to synaptic vulnerability, circuit instability and cognitive deficits in a subset of patients with SCZ (5,71). Moreover, the clinical relevance of interferon dysregulation is underscored by evidence that exogenous interferon exposure induces depressive and psychotic symptoms, further supporting a causal role for excessive interferon signaling in neuropsychiatric pathology (72). Collectively, our findings identify astrocytic IFN-β as a GABA-sensitive immune axis and suggest that GABA_A_ receptor activation via muscimol may exert neuroprotective effects by stabilizing astrocyte neuroimmune function and preventing interferon-driven maladaptive synaptic remodeling. These results provide mechanistic support for a model in which astrocyte-intrinsic immune modulation represents a key therapeutic target for limiting neuroinflammation-associated circuit dysfunction in SCZ.

Human clinical and translational studies demonstrate that muscimol reaches sub-micromolar to micromolar concentrations within the human CNS. Direct brain infusion studies reported venous plasma concentrations of ∼0.4 µM and CSF concentrations of ∼4 µM, with pharmacodynamic modeling indicating neuronal effects at tissue concentrations as low as ∼150 nM (73). In addition, a phase 1 clinical trial using intracerebral muscimol delivery employed infusion paradigms producing micromolar concentrations in human brain tissue, and human serum measurements following exposure have demonstrated concentrations approaching ∼1 µM (74,75). Collectively, these findings justify the concentration used in our in vitro systems, and support that 1 µM muscimol represents a therapeutically relevant concentration range for human CNS exposure for future clinical trials.

Several limitations of this study should be considered. First, although hCSs and iPSC-derived astrocytes provide physiologically relevant models that recapitulate key aspects of human neurodevelopment and cell-type-specific function, they cannot fully reproduce the cellular diversity, cytoarchitecture, and circuit-level complexity of the intact human brain, which might have influenced the repertoire of inflammatoty and plasticity responses we examined in this study. Second, the relatively modest number of donor-derived lines, while typical for iPSC-based mechanistic studies, may limit statistical power and the ability to capture the full spectrum of inter-individual variability inherent to SCZ. Finally, although integration with human postmortem transcriptomic datasets strengthens the translational relevance of our findings, direct causal relationships between GABA_A_ receptor-mediated immune modulation, astroglial functional rescue, and clinical phenotypes will require validation in animal models and human clinical studies. Despite these limitations, the convergence of transcriptomic, proteomic, and functional evidence across complementary experimental systems supports the robustness and translational relevance of the identified mechanisms.

Taken together, our findings support a model in which SCZ-associated immune dysregulation, particularly interferon-mediated antiviral signaling, disrupts astrocyte function and synaptic homeostasis, thereby impairing neuroplasticity. Activation of GABA_A_ receptors by muscimol reverses these abnormalities through suppression of inflammatory signaling and the simultaneous induction of neuroplastic pathways. These results establish non-classic psychedelic compounds as modulators of neuroimmune-plasticity coupling and identify astrocytes as a potential drug target and key cellular mediator linking immune dysfunction and synaptic pathology in SCZ.

## ACKNOWLEDGEMENTS

The research that led to this review received funding from the Research Council of Norway (Grant No. 223273), EU’s Horizon Psych-STRATA project (Grant No. 101057454), and the KG Jebsen Stiftelsen.

